# M_5_ positive allosteric modulation alleviates parkinsonian motor deficits

**DOI:** 10.1101/2025.08.21.671539

**Authors:** Nicole E. Chambers, Dominic Hall, Morgan Kaplan, Sarah Garan, Tyler Eichner, Mark S. Moehle

**Author notes:** Corresponding Author: Dr. Mark S. Moehle, Department of Pharmacology & Therapeutics University of Florida, 1200 Newell Drive, ARB R5-183 Gainesville, FL 32610.

## Abstract

Parkinson’s disease is a neurodegenerative movement disorder which is characterized by cardinal motor symptoms of tremor at rest, rigidity, bradykineasia, and postural instability. Underlying these cardinal motor symptoms is thought to be death and dysfunction of nigrostriatal dopamine neurons, and the gold-standard treatment of Parkinson’s disease is dopamine replacement therapy with the dopamine precursor L-DOPA. While efficacious, L-DOPA does not treat all motor symptoms and can have serious treatment-related side effects called L-DOPA induced dyskinesias, indicating an immense need for new targets to modulate dopaminergic function for anti-parkinsonian efficacy. One such potential target is the M_5_ muscarinic acetylcholine receptor, which has a unique expression profile where it is selectively expressed in midbrain dopaminergic neurons and their terminals in the striatum, and previous studies have indicated that M_5_ can modulate dopamine release and patterning of firing of dopamine neurons. Given this unique expression profile and function of M_5_, this receptor has an untested potential to modulate parkinsonian motor phenotypes. To test the potential for M_5_ to modulate Parkinsonian-like motor deficits and dyskinesia, we employed the unilateral 6-OHDA lesioned mouse model to create a hemi-parkinsonian state. Using multiple behavioral assays, including the cylinder test, forepaw adjusting steps assay, and in the Erasmus ladder, in conjunction with prototypical M_5_ pharmacological tool compounds, we investigated the ability of M_5_ to modulate parkinsonian motor deficits. Additionally, we tested the ability of M_5_ to modulate established L-DOPA induced dyskinesia or cause dyskinesia on its own. Overall, we found that M_5_ PAM alleviates forepaw asymmetry, bradykinesia, and spatial aspects of gait in the Erasmus ladder. Excitingly, M_5_ PAM does not cause robust dyskinesia, does not affect already established L-DOPA-induced dyskinesia, and does not affect L-DOPA motor efficacy. Taken together with previous findings, the current study suggests that M_5_ receptors are an exciting novel therapeutic strategy for ameliorating parkinsonian motor deficits even in late-stage models of severe PD without lessening L-DOPA’s motor benefit and without affecting existing symptoms of L-DOPA-induced dyskinesia.

## Introduction

Parkinson’s disease is a neurodegenerative movement disorder classically characterized by death of nigrostriatal dopamine (DA) neurons, and the development of cardinal motor symptoms(1). Standard treatment for PD is DA replacement therapy with L-DOPA(2). However, chronic L-DOPA treatment is associated with side effects including the development of motor fluctuations, such as “on” and “off” states, in up to 84% of PD patients(3), and up to 20% of patients experience gait symptoms which are not improved by L-DOPA(4). Additionally, in up to 90% of PD patients(5), chronic L-DOPA treatment results in debilitating, abnormal involuntary movements which are called L-DOPA-induced dyskinesia. The lack of treatment of all motor symptoms by L-DOPA, the dosing problems, and the debilitating side effects of chronic L-DOPA treatment necessitate new PD therapeutics to meet a critical unmet need.

One possible therapeutic mechanism is through targeting acetylcholine receptors Multiple studies have highlighted acetylcholine (ACh) as a critical modulator of motor function and LID, with a critical role for muscarinic acetylcholine receptors in regulating neurons and circuits that are associated with these symptoms (6–11). The muscarinic acetylcholine receptors are group of 5 metabotropic receptors termed M_1_ through M_5_ (12). Of these, M_1_ and M_4_ are robustly expressed in basal ganglia nuclei, and have had functions in normal and pathological states that are prescribed as regulating parkinsonian phenotypes(6, 13–16). Additionally, anti-muscarinic drugs (namely trihexyphenidyl) are efficacious at removing the primary motor symptoms of PD(17). Recent studies have even suggested that the efficacy of these anti-muscarinic therapeutics is through blocking the actions of acetylcholine specifically on the M_4_ receptor (15, 18–20). However, the M5 receptor may present a unique opportunity to modulate SNc dopamine neurons for relief of parkinsonian motor symptoms. The Gq-coupled M_5_ muscarinic acetylcholine receptor (M_5_) may be a particularly promising therapeutic target for PD due to its exclusive location on DA neuron cell bodies in the substantia nigra and ventral tegmental area, as well as nigrostriatal DA terminals (21, 22). Activation of M_5_ can modulate nigrostriatal DA release and dopamine neuron firing rate and patterning (23–25), Conversely blocking M_5_ can inhibit the rewarding effects of drugs of abuse, and co-formulation of M_5_ negative allosteric modulators with drugs of abuse has been suggested as a mechanism to block the addictive properties of opioids while maintaining the anti-nociceptive efficacy of these compounds(25–28). Overall, the selective expression of M_5_ coupled with proposed actions of M_5_ on midbrain dopamine neurons, suggests that M_5_ positive allosteric modulators or activators could potentially influence PD motor symptoms and LID. However, this idea is completely untested in relevant parkinsonian models.

Here, we utilize the unilateral 6-OHDA lesioned mouse model of severe Parkinsonian-like motor deficits coupled with gold-standard prototypical M_5_ positive or negative allosteric modulators to directly test this idea. This study investigated the role M_5_ allosteric modulators on motor deficits and LID through robust, well described tests of parkinsonian motor deficits and dyskinesia as well as through novel automated tests of motor performance. Overall, our results suggest that M_5_ positive allosteric modulators improves motor and gait metrics but not overall akinesia in the 6-OHDA lesion model of PD. Furthermore, our results show that M_5_ PAM does not significantly worsen LID or cause dyskinesia on its own. Taken together, these results provide the first evidence to indicate a potential therapeutic benefit for M5 receptor modulation to relieve specific parkinsonian motor deficits.

## Materials and methods

### Subjects

Male C57BL6/J mice were used in these experiments, n = 32. A total of 22 mice underwent 6-OHDA lesion surgery (n = 22), and naïve age-matched controls were used as comparators for the Erasmus Ladder experiment examining 6-OHDA-induced motor deficits, (n = 10). Mice were housed in a colony room kept at a constant temperature with a 12 h light and dark cycle (lights on at 06:00), in cages of up to 5 mice. These mice were given *ad libitum* food and water. All procedures executed were in accordance with and approved by the University of Florida Institutional Animal Care and Use Committee.

### Surgery

One week prior to surgery mice were habituated to Stat caloric supplement to stave off weight loss from surgery. Similar to Chambers et al., (29, 30) mice were anesthetized with 2-3% isoflurane in oxygen, injected with carprofen (15 mg/kg) as a preoperative analgesic, desipramine was injected to prevent the loss of norepinephrine neurons, bupivacaine was injected under the scalp (0.2 mL) 5 min prior to the incision. Thereafter animals were stabilized in an RWD stereotax and three alternating saline and chlorhexidine scrubs were applied prior to the incision. Next, a burr hole was drilled at the target coordinates (AP: -1.0 mm, ML: -1.0 mm, DV: -4.9 mm) over the right medial forebrain bundle and 6-OHDA (6-hydroxydopamine hydrobromide, Sigma) was injected at a rate of 0.1 uL/min, at a concentration of 7.5 µg / µL (free base), for a total volume of 1 µL. 6-OHDA was formulated in a vehicle of 0.2% ascorbate in sterile saline. After the injection, the needle was left *in situ* for 5 min and then slowly withdrawn. The scalp was closed with sterile sutures and Vetbond. Mice were given 0.3 mL of saline (s.c.), moistened chow, and stat caloric supplement to stave off weight loss and dehydration. Additionally, meloxicam (15 mg/kg) was given as a postoperative analgesic every 24 h for 48 h following surgery.

### Experimental Design

All mice other than age-matched controls for Erasmus ladder underwent 6-OHDA lesion surgery (See Fig 1A for Experimental Timeline and Design). Two cohorts of 6-OHDA animals were used in the experiments for this paper, n = 22. All animals underwent baseline (off treatment) testing with the cylinder test of forelimb asymmetry to verify 6- OHDA lesion. Thereafter, in a within-subjects counterbalanced design all mice were ran through 3 test days for baseline (off treatment), 30 mg/kg of NAM, and 100 mg/kg PAM while testing forehand adjusting steps, Erasmus ladder, and ALO AIMs 60 min after treatment. Critically, at least 2 days of drug washout were provided between test days. A control age-matched cohort of male mice (n = 10) ran through the Erasmus ladder off-treatment to determine differences between 6-OHDA lesioned mice and control mice.

**Figure 1.**
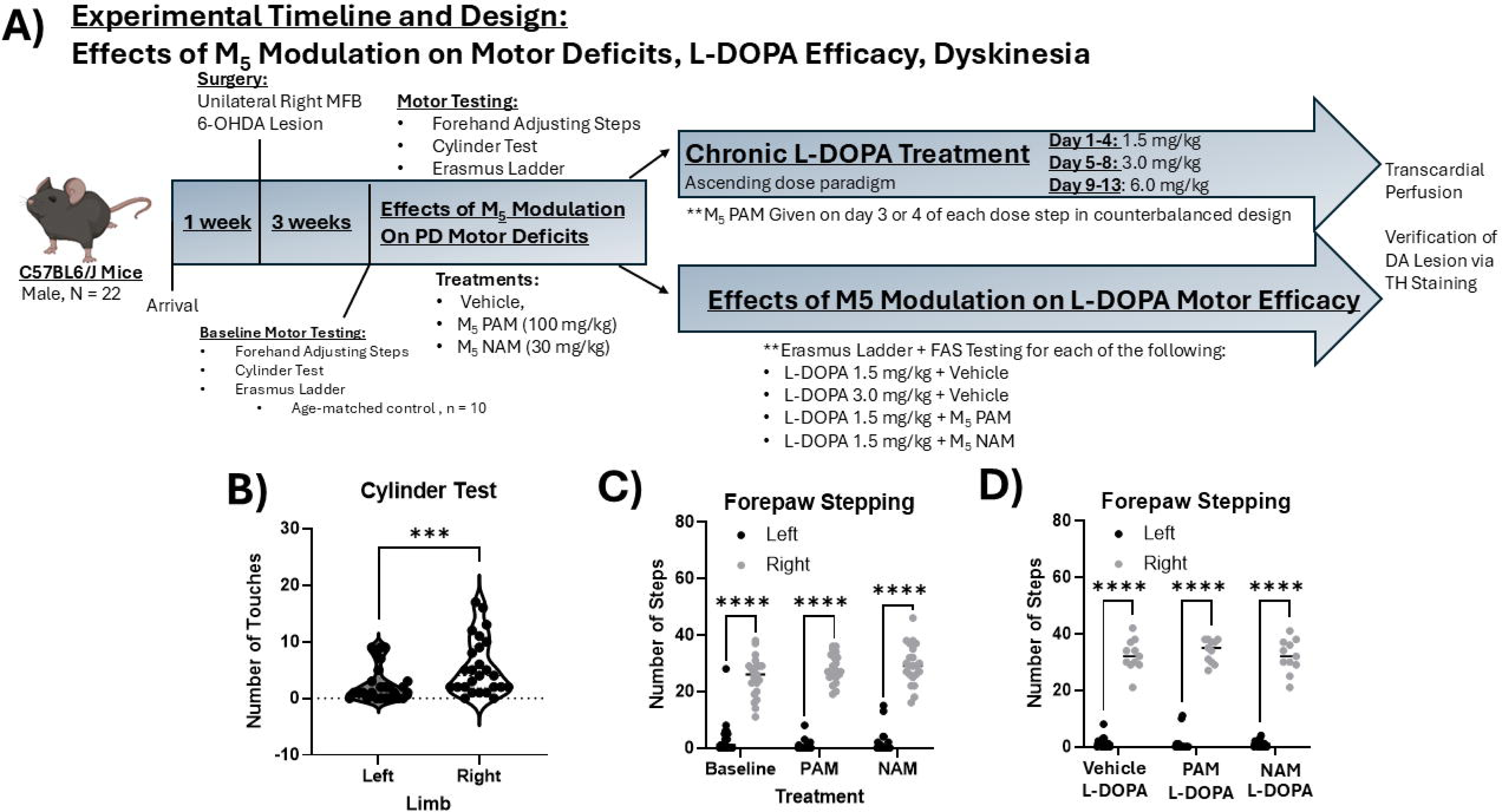
Experimental Timeline and Effects of M_5_ Modulation on Parkinsonian Forepaw Deficits. A) A schematic showing the timeline and design of this study. A total of 22 C57BL6/J mice were included in this study, they arrived and were habituated to STAT caloric supplement and mush food daily for one week before surgery. Following this, a unilateral right medial forebrain bundle (MFB) 6-OHDA lesion was performed. Mice were given 3 weeks for recovery, during which they were given STAT caloric supplement and mush food daily. At three weeks after surgery mice underwent baseline motor testing with cylinder test and an adapted version of the forehand adjusted steps test for mice to confirm lesion. Unlesioned mice were excluded from further study (forehand percent intact score above 40%). Mice then underwent a battery of motor testing to study the effects of M5 modulation on PD motor deficits which included cylinder test, forepaw adjusting steps test, and Erasmus ladder testing. In a within-subjects, counterbalanced design, mice were M5 PAM, VU0238429 (100 mg/kg, i.p.), or M5 NAM ML375 (30 mg/kg, i.p.) to understand their effects on baseline motor deficits resulting from lesion. Thereafter, mice were divided into subcohorts for further participation in a study on M5 modulation effects on L-DOPA motor efficacy and M5 modulation effects on L-DOPA-induced dyskinesia. In the L-DOPA-induced dyskinesia experiment, mice were given daily, chronic L-DOPA treatment in an ascending dose paradigm where each dose was given on 4 days, (1.5, 3.0, and 6.0 mg/kg). For each dose step on the 3^rd^ and 4^th^ day, in a within-subjects, counterbalanced design mice received vehicle + L-DOPA or M_5_ PAM + L-DOPA. L-DOPA-induced dyskinesia was scored using the validated rodent Abnormal involuntary movements scale (AIMs) beginning 20 min after injection. Mice were video recorded in a 3-D neurobehavioral apparatus designed to show all body angles, and data were later scored using the AIMs scale for 1 min every 20 min for 140 min. For the effects of M5 modulation on L-DOPA efficacy, we again used cylinder test, forepaw adjusting steps, and the Erasmus ladder to understand how M5 modulation affected L-DOPA-mediated behavior. In a within-subjects, counterbalanced design, mice were given 1.5 mg/kg L-DOPA, 3.0 mg/kg L- DOPA, L-DOPA(1.5 mg/kg) + M5 PAM, and L-DOPA(1.5 mg/kg) + M5 NAM. All behavior for these experiments was run 1 hour after injection of L-DOPA and PAM or NAM, as this corresponds to peak brain plasma expression of both L-DOPA and the M5 compounds. B) Shows forepaw asymmetry in the cylinder test resulting from 6-OHDA lesion at baseline. Data were analyzed via paired-samples t-test. *** p < 0.001. C) Shows forepaw stepping from forehand adjusting steps test. Data were analyzed via two-way repeated-measures ANOVA with within-subject factors of treatment (3: Baseline (off treatment), PAM, NAM), and side (2: left(lesioned) paw, right (intact) paw). **** p < 0.0001. Overall there was no effect of treatment on the akinetic motor deficit in the left (lesioned) forepaw. D) Shows forepaw stepping from forehand adjusting steps test on vehicle + L-DOPA, PAM (100 mg/kg) + L-DOPA, or NAM (30 mg/kg) + L-DOPA. As in C, a two-way repeated measures ANOVA was conducted, which revealed that there were no effects of treatment but there was an effect of lesion, with the left (lesioned) forepaw taking significantly less steps despite treatment. **** p < 0.0001.

Thereafter, the two 6-OHDA lesioned cohorts were divided for further testing with L-DOPA. Cohort 1 was employed to understand the effects of M_5_ PAM or NAM on L-DOPA-mediated motor improvement. In a within-subjects, counterbalanced design they were given L-DOPA (1.5 mg/kg) + vehicle, L-DOPA (1.5 mg/kg) + PAM, and L-DOPA (1.5 mg/kg) + NAM. Cohort 2 was employed in an ascending dose study to understand the effects of M_5_ PAM on L-DOPA-induced dyskinesia. Each dose step (1.5, 3, and 6 mg/kg) lasted for 4 days. On the third and fourth day of each dose in a within-subjects, counterbalanced design, mice in cohort 2 received either L-DOPA + vehicle or L-DOPA + M_5_ PAM (100 mg/kg). All unlesioned mice, as shown by <40% reduction in cylinder test asymmetry and forepaw adjusting steps, were excluded from the study.

### Pharmaceutical treatment with M_5_ PAM and L-DOPA

M_5_ negative allosteric modulator ML375 (30 mg/kg, i.p.) was used to test the effects of M_5_ inhibition on behavior(31). M_5_ positive allosteric modulator VU0238429 (100 mg/kg, i.p.) was used to test the effects of M_5_ potentiation on behavior (32). Vehicle for both M_5_ PAM and NAM was 10% Tween 80 dissolved in sterile water containing 20% hydroxypropyl Beta-cyclodextran. When motor behavior was tested, these compounds were given 60 min before behavior as this corresponds with peak brain plasma levels. L-DOPA was injected at doses of 1.5, 3, or 6 mg/kg (i.p). Importantly, L-DOPA was always given with the peripheral decarboxylase inhibitor Benserazide hydrochloride (15 mg/kg). On days when both L-DOPA and M_5_ PAM were administered, they were injected at the same time, one hour before motor behavior testing other than abnormal involuntary movements testing, which began 20 min after injection.

### Behavioral Testing

#### Cylinder test

To quantify motor deficits arising from 6-OHDA lesion and to validate our adaptation of forepaw adjusting steps for mice, we used the cylinder test of forepaw asymmetry (33–35). We placed mice into a glass cylinder for 10 min while video recording to measure paw touches on each side. Paw touches were evaluated based on which paw touched first.

#### Forepaw adjusting steps (FAS)

To quantify motor deficits arising from our 6-OHDA lesion and motor effects of L-DOPA and M_5_ modulators, we adapted methods from Chang et al., (36) to mice. Mice were gently scruffed and were dragged across the counter such that only one of their forepaws touched the counter. Mice were dragged laterally over a table for a length of 30 cm over 10s. Forepaw adjusting steps were counted by a trained rater who was blind to condition. Forepaw percent intact scores were calculated by taking steps with the left (lesioned) forepaw divided by the right (unlesioned) forepaw and multiplying the quotient by 100.

#### Erasmus ladder

To evaluate motor performance, motor coordination, and gait, the Erasmus ladder was used (37, 38). Briefly, mice walk along a ladder which consists of high and low rungs and their steps are quantified by on-board computers for 42 trials for each mouse. Data presented represent the average of 42 trials for baseline (off-treatment) and on-treatment conditions.

#### L-DOPA-induced dyskinesia

On test days, mice were habituated for 15 min to glass cylinders before being injected with L-DOPA. Thereafter a behavioral video was recorded in a custom 3-D Neurobehavioral Chamber for 1 min every 20 min for 120 min to later quantify LID. Videos were scored using the validated abnormal involuntary movements scale (AIMS(39–42)) as described in (30).

### Statistical analyses

All statistical analyses were performed in SPSS with alpha set to 0.05. Cylinder test data were analyzed using a paired-samples t-test comparing right (intact) vs. left (lesioned) limb. For comparisons between multiple groups for FAS and Erasmus ladder, parametric repeated-measures one-way or two-way ANOVAs were used with Tukey post-hoc analysis as appropriate. In the Erasmus ladder, in cases where a particular type of step was only taken by a subset of animals, a mixed-effects analysis was used with Dunnett’s post-hoc tests. For LID behavior and comparisons of ALO scores for peak plasma concentrations of M5 PAM vs. L-DOPA, we used Friedman ANOVA with Dunn posthocs was used to compare groups.

## Results

### Hemi-parkinsonian mice show forepaw akinesia which is not improved by M_5_ modulation

The cylinder test was conducted to confirm lesion using measurements of forelimb asymmetry in hemi-parkinsonian mice (Fig 1B). A paired-samples t-test revealed that there were significantly less cylinder touches with the left (lesioned) forepaw than for the right (unlesioned) forepaw, t(25) = 2.75, p = 0.0108, indicating that mice had significant nigral dopamine lesions.

To assess akinesia, we adapted the forepaw adjusting steps test to mice(36) to assess the effects of M_5_ modulators and L-DOPA on akinesia (Fig 1C-D). In M_5_ modulator treated mice, we found a significant difference between performance of left and right paws in this assay, but this was not ameliorated by treatment with M_5_ PAM or NAM (Figure 1C, two-way repeated measures ANOVA, main effect F(2,84 = 4.36, p = 0.016, side effect F(1, 42) = 559.6, p = 0.0001). Overall, this shows the lack of an effect of M_5_ modulation to directly improve forepaw stepping as a measure of akinesia.

Similarly, we tested whether M_5_ PAM or NAM could potentiate sub-threshold doses of L-DOPA (1.5 mg/kg) to ameliorate akinetic phenotypes. We did not observe M_5_ PAM or NAM being able to modulate forepaw adjusting steps in the presence of L-DOPA (Figure 1D). Taken together, this suggests that M_5_ modulators cannot improve severe akinetic phenotypes in the hemi-parkinsonian mouse either directly or in combination with L-DOPA treatment.

### Hemi-parkinsonian mice show bradykinesia and short steps in Erasmus ladder

To sensitively measure parkinsonian motor deficits in an unbiased manner across multiple measures, we utilized an automated Erasmus Ladder to assess motor performance and gait in 6-OHDA lesioned mice and controls. We found differences resulting from the 6-OHDA lesion across multiple metrics in the Erasmus ladder that indicated that the automated Erasmus Ladder could sensitively and robustly measure parkinsonian motor phenotypes (Figure 2). Although overall trial duration was not affected (Fig 2A) hemi-parkinsonian mice took longer to complete a variety of steps as compared to age matched controls (see Table 1 for statistical values, whether Welch’s correction was used, and interpretations). Specifically, 6-OHDA lesioned mice took longer to complete high rung backsteps t(23.91) = -2.92, p = 0.007 (Fig 2B), high rung short steps t(25.76) = -2.34, p = 0.014 (Fig 2C), high rung long steps t(21.46) = -4.52, p < 0.0001 (Fig 2D), high rung jumps t(21.45) = -2.94, p = 0.012 (Fig 2E), high to low rung short steps t(24.35) = -2.94, p =0.007 (Fig 2F), high to low rung long steps, t(21.99) = - 3.39, p = 0.003 (2G), low to high rung long steps t(25.51) = -3.55, p = 0.002 (Fig 2H), and low to high rung jumps t(25.75) = -2.36, p = 0.026 (Fig 2I). This indicates a bradykinesia phenotype in our 6-OHDA lesioned mice. We also observed that 6-OHDA lesioned mice take more high rung short steps t(30) = -2.58, p = 0.008 (Fig 2J), and less high rung long steps t(21.07) = 3.46, p = 0.001 than age-matched controls, consistent with the short, shuffling steps seen in human PD patients. See Table 1 for statistical values.

**Figure 2.**
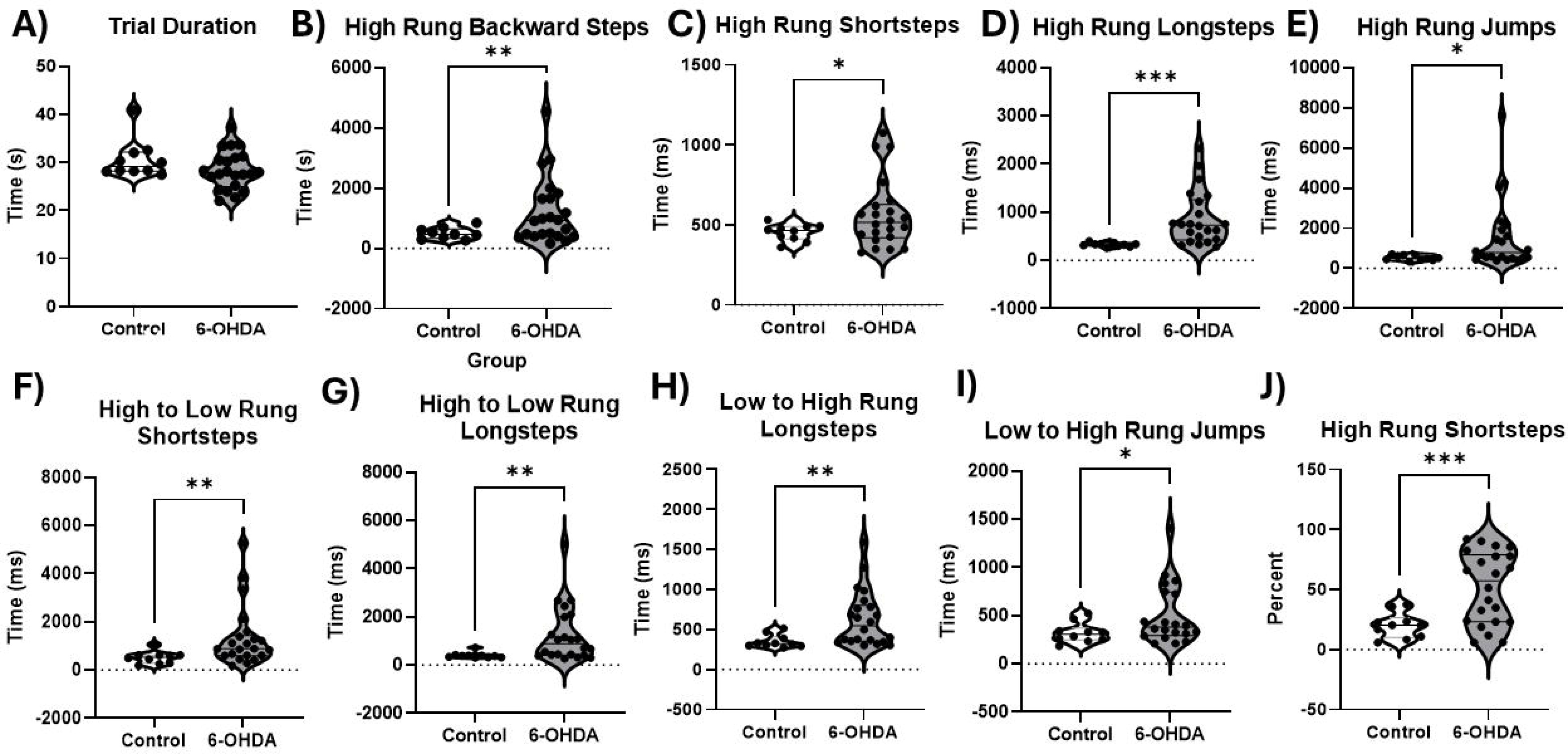
Differences Between Age-Matched Control and 6-OHDA Lesioned Mice in Erasmus Ladder. For all data included in this section, we used age-matched C57BL6/J mice (n = 10) as comparators to our 6-OHDA lesioned C67BL6/J mice (n = 22). This is the first description of Eramsus ladder deficits in 6-OHDA lesioned mice. All data were analyzed using two-tailed independent-samples t-tests, alpha = 0.05. Overall, we observed that 6-OHDA lesioned mice show bradykinesia in multiple aspects of the Erasmus ladder, A –I. A) There is no change in overall trial duration between 6-OHDA lesioned and control mice. B) 6-OHDA lesioned mice take more time to complete backward steps on the high rung. C) 6-OHDA lesioned mice take more short steps on the high rung than controls, which recapitulates aspects of human gait in Parkinson’s disease, as human PD patients show short, shuffling steps. D) 6-OHDA lesioned mice take more time to complete long steps on the high rung. E) 6-OHDA lesioned mice take more time to complete long steps on the high rung. E) 6-OHDA lesioned mice take longer to make high rung jumps than controls, F) 6-OHDA lesioned mice take longer to complete High to low rung shortsteps G) 6-OHDA lesioned mice take longer to complete high to low rung longsteps, H) 6-OHDA lesioned mice take longer to complete low to high rung longsteps. I) 6-OHDA lesioned mice take longer to complete low to high rung jumps. J) 6-OHDA lesioned mice take a higher percentage of high rung short steps than control mice, showing a clear parkinsonian phenotype. * p < 0.05, ** p < 0.01, *** p < 0.001

**Table 1.** Erasmus Ladder Deficits in 6-OHDA Lesioned Mice vs. Controls.

| Rung | Variable | df | t | p<sup>a</sup> | Equal Variance | Interpretation |
| --- | --- | --- | --- | --- | --- | --- |
| High rung backsteps | Time | (23.91) | -2.92 | 0.007 | No | 6-OHDA slower than control |
| High rung shortsteps | Time | (25.76) | -2.34 | 0.014 | No | 6-OHDA slower than control |
| High rung longsteps | Time | (21.46) | -4.52 | < 0.0001 | No | 6-OHDA slower than control |
| High rung jumps | Time | (21.45) | -2.94 | 0.012 | No | 6-OHDA slower than control |
| High to low rung shortsteps | Time | (24.35) | -2.94 | 0.007 | No | 6-OHDA slower than control |
| High to low rung longsteps | Time | (21.99) | -3.39 | 0.003 | No | 6-OHDA slower than control |
| Low to high rung long steps | Time | 25.51 | -3.55 | 0.002 | No | 6-OHDA slower than control |
| Low to high rung jumps | Time | 25.75 | -2.36 | 0.026 | No | 6-OHDA slower than control |
| High rung shortsteps | Percent | 30 | -2.58 | 0.008 |  | 6-OHDA takes more short steps than control |
| High rung longsteps | Percent | 21.07 | 3.46 | 0.001 | No | 6-OHDA takes less long steps than control |

df degrees of freedom, 6-OHDA 6-hydroxydopamine
<sup>a</sup> p values were calculated using a parametric independent samples t-test

### Treatment with M_5_ PAM Alleviates Erasmus Ladder Deficits

To assess anti-parkinsonian motor efficacy of M_5_ receptor PAM VU0238429 and M_5_ NAM ML375, we injected these M_5_ modulators 1 hour before performing behavioral testing, as this corresponds to peak brain plasma concentrations. Unless otherwise stated, data were analyzed using a one-way repeated-measures ANOVA with within-subjects factor of drug treatment (3: baseline off treatment, M_5_ PAM, M_5_ NAM).

However, not all mice take certain types of steps in the Erasmus ladder (e.g. backward steps). In these cases we have used a mixed-effects analysis. In either case, Dunnett’s multiple comparisons test was used as appropriate to clarify significant results between baseline and drug conditions. Overall, we observed that the M_5_ PAM could improve performance in multiple aspects of Erasmus ladder performance in 6-OHDA lesioned mice, detailed below (see Table 2 for statistical values, Fig 3).

**Figure 3.**
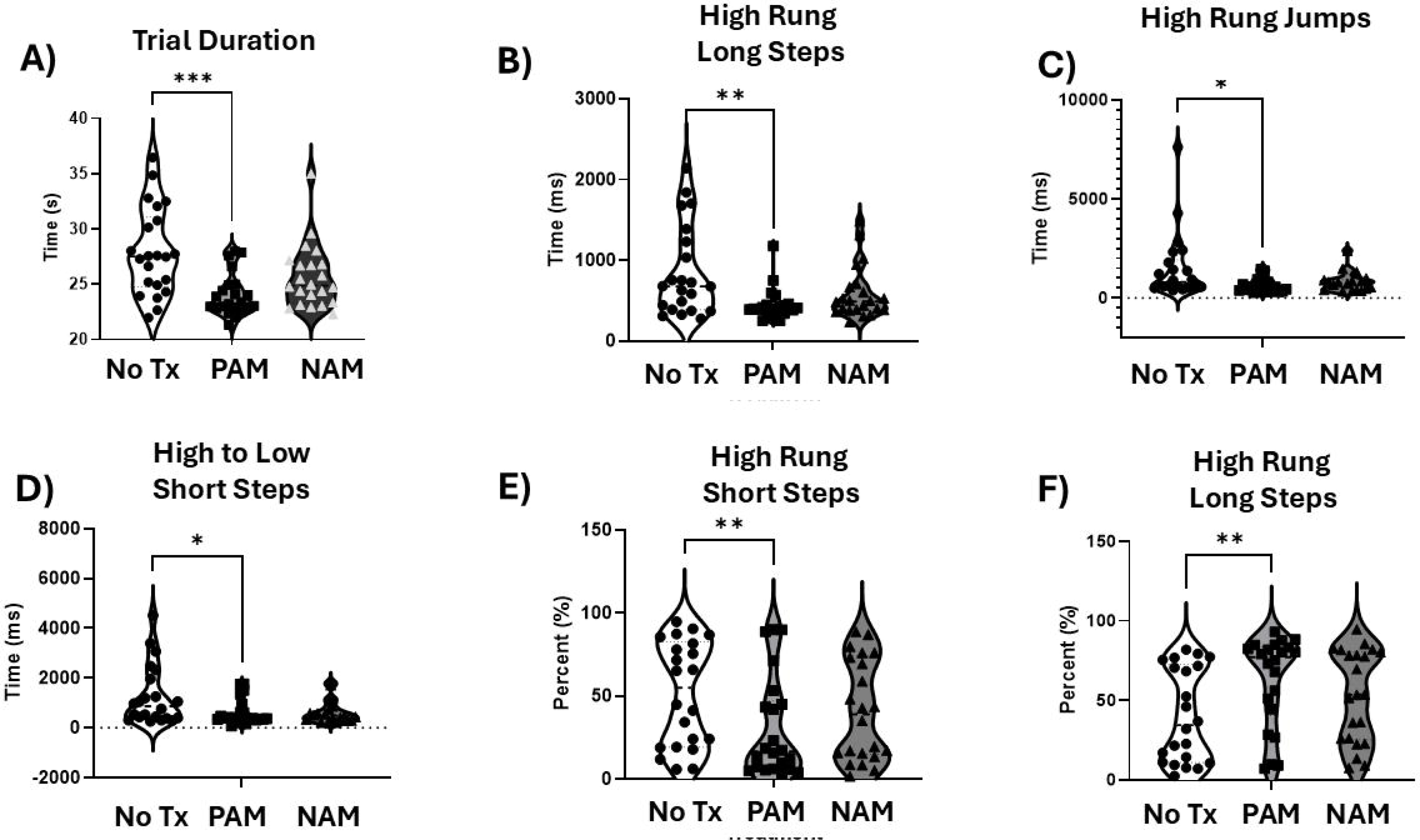
M5 Receptor Modulation Alleviates Erasmus Ladder Deficits in 6-OHDA Lesioned Mice. All Erasmus ladder data were analyzed using a repeated-measures ANOVA with factor of treatment (3: no treatment baseline, PAM, NAM). Dunnett tests were used as necessary (PAM and NAM treatment were only compared with Baseline). Mice were injected with M5 PAM VU0238429 (100 mg/kg, i.p.) or M5 NAM ML375 (30 mg/kg, i.p.) one hour before behavioral testing. A) There was a decrease in overall trial duration for mice treated with M5 PAM, suggesting the M5 PAM alleviates bradykinesia in the Erasmus ladder, *** p < 0.001 B) Mice treated witih M5 PAM take less time to complete high rung long steps C) Mice treated with M5 PAM show a reduction in the time it takes to complete high rung jumps. D) Mice treated with M5 PAM take less time to complete high to low rung short steps E) Mice treated with M5 PAM take a smaller percentage of short steps on the high rung, suggesting that M5 PAM improves stride length F) Mice treated with M5 PAM show a higher percentage of high rung long steps. Overall data in this figure suggest that M5 PAM alleviates Parkinsonian-like deficits in stride length and bradykinesia in our 6-OHDA lesioned model. * p < 0.05, ** p < 0.01, *** p < 0.001

**Table 2.** Erasmus Ladder Motor Improvements with M_5_ PAM and NAM.

| Rung/Variable | Variable<br>Type | df | F | p | Test | Interpretation |
| --- | --- | --- | --- | --- | --- | --- |
| Trial Duration | Time | 1.681,<br>35.29 | 9.592 | 0.0009,<br>0.0001<br>PAM vs.<br>Off-Tx | Mice took<br>less time to<br>complete<br>trials with<br>PAM | Mice took less<br>time to complete<br>trials with PAM |
| High Rung Long<br>steps | Time | 1.592,<br>33.43 | 6.481 | 0.0070 | Mice with<br>PAM take<br>less time to<br>complete<br>high rung<br>longsteps<br>0.0052<br>Baseline vs.<br>PAM | Mice with PAM<br>take less time to<br>complete high<br>rung longsteps<br>0.0052 Baseline<br>vs. PAM |
| High Rung<br>Jumps | Time | 1.160,<br>20.87 | 5.337 | 0.027 | Mixed<br>Effects<br>Analysis | Mice with PAM<br>take less time to<br>complete high rung jumps than<br>at baseline, p =<br>0.033 |
| High to Low<br>Rung Short<br>Steps | Time | 1.473,<br>29.45 | 5.86 | 0.013 | Mixed<br>Effects<br>Analysis |  |
| High Rung<br>Short Steps | Percent | 1.869,<br>39.24 | 3.728 | 0.0356,<br>0.0086 | Mixed<br>Effects<br>Analysis | Mice with PAM<br>take less short<br>steps than with<br>baseline |
| High Rung Long<br>Steps | Percent | 1.753,<br>36.82 | 4.306 | 0.025 |  | Mice with PAM<br>take more long<br>steps than<br>baseline |

df degrees of freedom, PAM positive allosteric modulator
<sup>a</sup> p values were calculated using a one-way repeated measures ANOVA with Tukey posthoc testing as appropriate

There were effects of treatment on overall trial duration F(1.681, 35.29) = 9.59, p = 0.0009 (Fig 3A), time to complete high rung long steps F(1.592, 33.43) = 6.48, p = 0.007 (Fig 3B), time to complete high rung jumps F(1.16, 20.87) = 5.337, p = 0.027 (Fig3C), and time to complete high to low rung short steps F(1.473, 29.45) = 5.86, p = 0.013 (Fig 3D). Overall, our analyses revealed that treatment with M_5_ PAM improved bradykinesia in the Erasmus ladder in our hemi-parkinsonian mouse model (see table 2 for statistical values). There were also effects of treatment on overall percentage of short steps taken on the high rung F(1.869, 39.24) = 3.728, p = 0.036 (Fig 3E), and on percentage of long steps taken on the high rung F(1.753, 36.82) = 4.306, p = 0.025 (Fig 3F). This suggests that M_5_ PAM improved spatial aspect of PD gait, reducing the number of short steps taken by hemi-parkinsonian mice and increasing the number of long steps. Overall, these data suggest that M_5_ PAM can reduce bradykinetic and gait dysfunction parkinsonian-like phenotypes in sensitive measures of motor performance and gait.

### L-DOPA alleviates certain Erasmus ladder, but does not act synergistically with M_5_ PAM

To understand if M_5_ modulators could synergistically interact with L-DOPA, we first ran mice through the Erasmus ladder one hour after L-DOPA (1.5 or 3 mg/kg), which corresponds to peak brain plasma levels of L-DOPA. We used these doses of L-DOPA because mice experienced LID and rotations that precluded the use of the Erasmus ladder at 6mg/kg of L-DOPA and higher. Overall, there was no effect of L-DOPA on overall trial duration (Fig 4A). However, mixed effects models revealed that L-DOPA (3.0 mg/kg) reduced the number of high rung short steps, p = 0.009 (Fig 4B), and resulted in a non-significant, but trending, reductions in time it took to complete high rung long steps (Fig 4C), and resulted in a non-significant, but trending, increases in percentage of high rung long steps (Fig 4D). We therefore used L-DOPA 1.5 mg/kg as the threshold dose to determine if M_5_ could potentiate its effects and act as an L-DOPA sparing strategy. We then co-injected L-DOPA (1.5 mg/kg) and M_5_ PAM or NAM on separate test days. One hour after co-injection, which corresponds to peak plasma levels of both L-DOPA and M_5_ compounds, we tested the effects of adding M_5_ PAM or NAM on L-DOPA in the Erasmus ladder. Overall, our results suggest that both M_5_ PAM and NAM do not potentiate L-DOPA’s motor benefit, as it does not significantly affect trial duration, high rung short steps, high rung long steps of percentage of high rung long steps (Fig 4E-H). These data suggest that efficacy of M_5_ PAMs observed above act in a distinct mechanism than that of L-DOPA.

**Figure 4.**
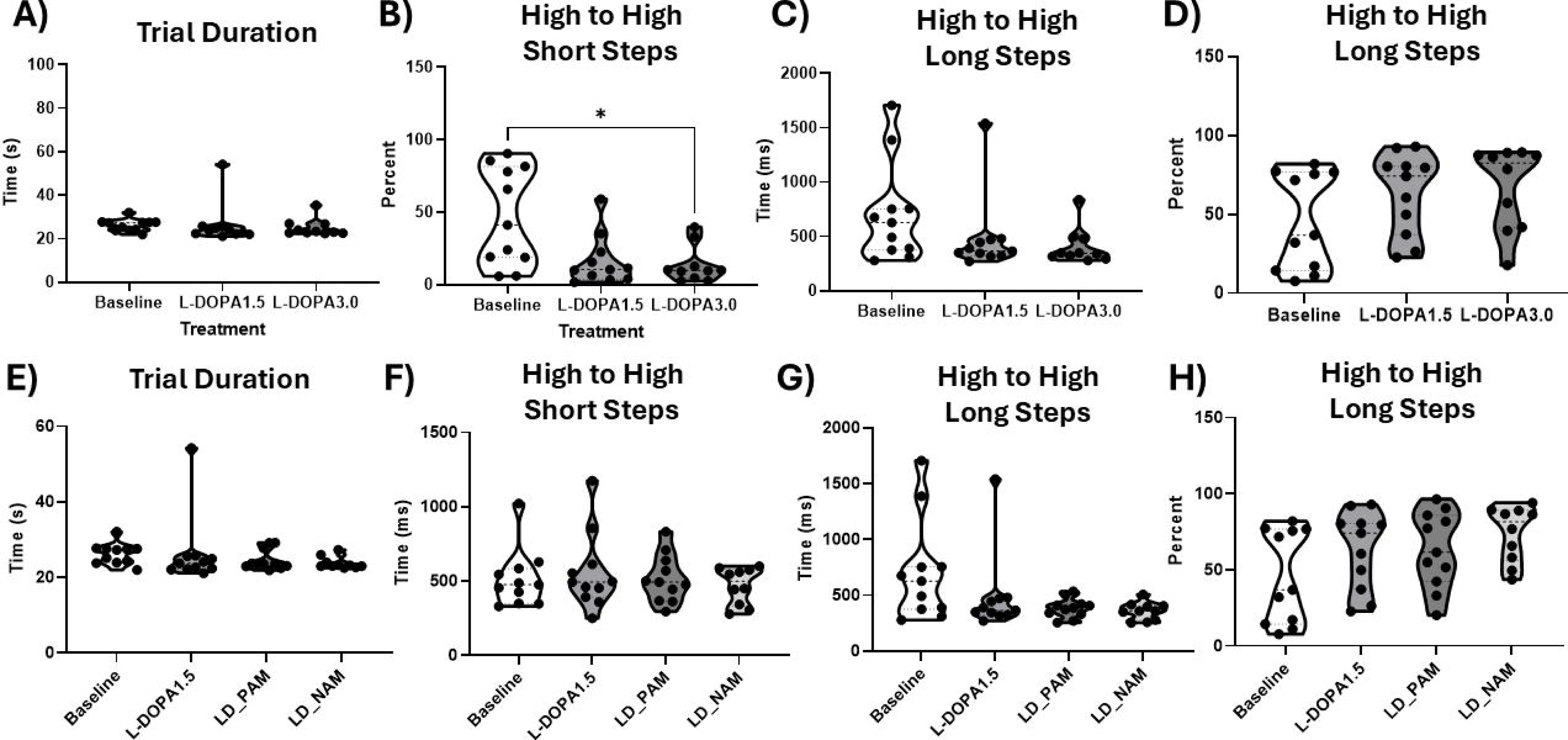
M_5_ PAM does not potentiate the effects of L-DOPA. All data were analyzed with a repeated-measures ANOVA with factor of treatment. Tukey post-hocs were conducted as appropriate to understand significant differences. Comparisons were only made between treatment groups and no treatment baseline A) Trial duration was not affected by L-DOPA. B) The percentage of high rung short steps was reduced by 3 mg/kg of L-DOPA, suggesting an improvement in stride length. C) Time to complete high rung long steps was not significantly reduced by L-DOPA. D) Percentage of high rung longsteps was aincreased in mice with 3.0 mg/kg of L-DOPA, again suugesting an improvement in stride length with L-DOPA. E) Overall trial duration was not affected when PAM or NAM were added to L-DOPA. F) Time to complete high rung short steps was not affected by co-treatment with PAM or NAM. G) Time to complete short steps was unaffected by L-DOPA co-treatment with M5 PAM and NAM. H) Percentage of high rung longsteps was not significantly changed by L-DOPA co-treatment with M5 PAM or NAM. Overall, this suggests that M5 PAM does not affect L-DOPA’s motor efficacy. * p < 0.05

### M_5_ PAM does not alter L-DOPA-induced dyskinesia

To understand the effects of M_5_ PAM on L-DOPA-induced dyskinesia, we injected L-DOPA in an ascending dose paradigm study (For experimental timeline and design see Fig 5A). In a within-subjects, counterbalanced design, we injected M_5_ PAM or vehicle on day 3 or 4 along with L-DOPA. For the ascending doses of L-DOPA, we used 1.5 mg/kg (Fig 5B), 3.0 mg/kg (Fig 5C), and 6.0 mg/kg (Fig 5D). We used a Kruskal Wallis test to compare L-DOPA alone vs. M_5_ PAM, with Dunn’s posthoc testing conducted as appropriate. Overall, we found that there were no significant differences in overall dyskinesia severity for 1.5, 3.0, or for 6.0 mg/kg of L-DOPA when paired with M_5_ PAM. This suggests that M_5_ PAM may be used for additional therapeutic benefit in PD without increasing motor side effects of L-DOPA. Furthermore, we determined whether M_5_ PAM alone could cause dyskinesia. Measuring LID using ALO AIMs during peak plasma concentration (60 min after injection), we found that M_5_ PAM did not elicit robust dyskinesia, with only some intermittent orolingual behavior observed. We compared the 60 min rating with the peak plasma concentration rating for L-DOPA 1.5, 3.0, and 6.0 mg/kg using a non-parametric Friedman ANOVA with Dunn’s post-hocs as appropriate F(4) = 21.27, p < 0.0001. Overall, we found that mice receiving M_5_ PAM alone did not show robust dyskinesia at peak plasma concentration of M_5_ PAM, which in contrast to the dyskinesia evident with peak plasma effects of 3 mg/kg of L-DOPA, p = 0.015 and 6 mg/kg of L-DOPA p = 0.0004 (Fig 5E).

**Figure 5.**
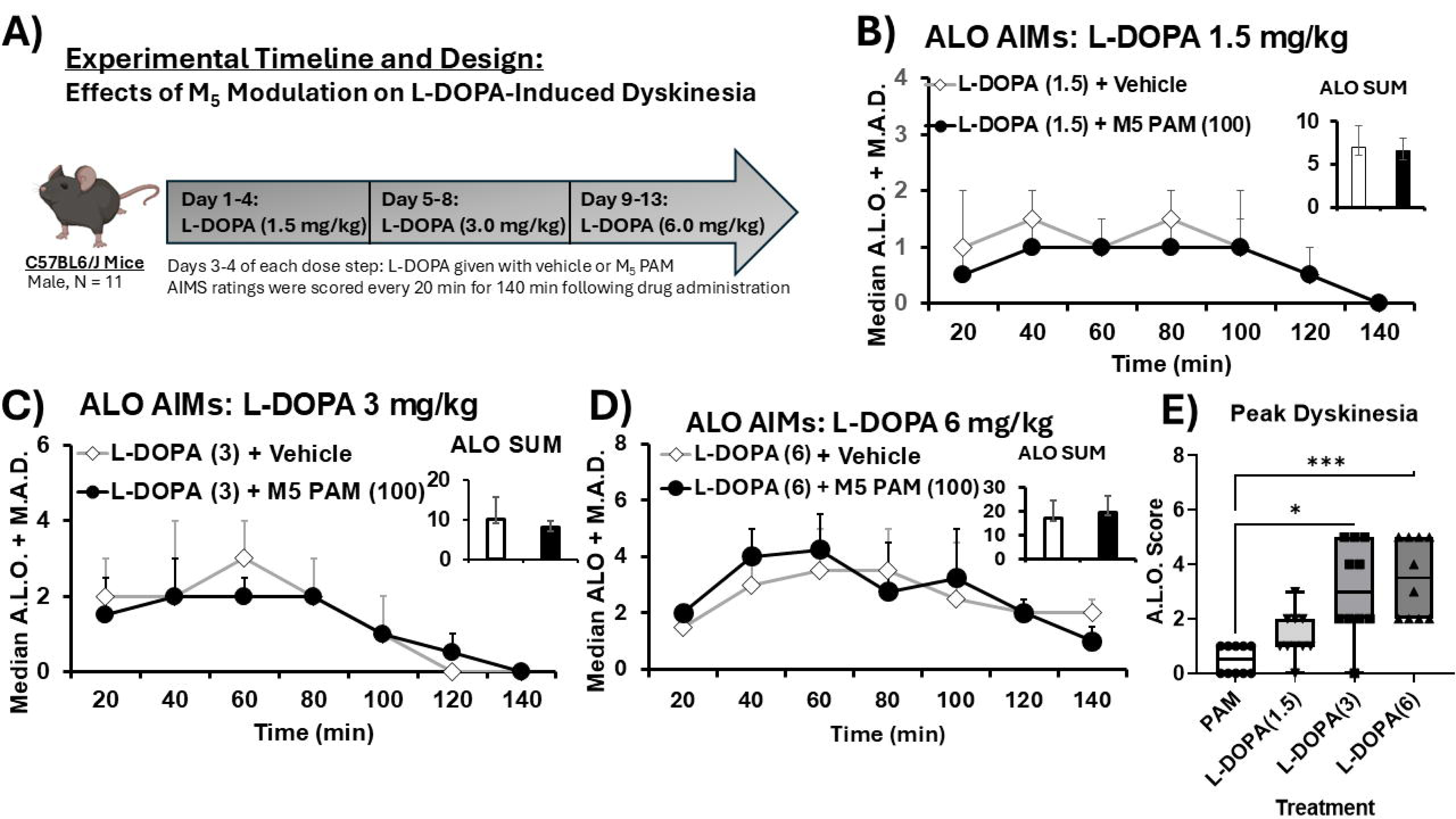
M_5_ Receptor Modulation Does Not Affect L-DOPA-Induced Dyskinesia And Does Not Cause Dyskinesia. Using a within-subjects design, mice were given chronic, daily L-DOPA treatment in an ascending dose paradigm (four days of each dose, 1.5, 3.0, and 6.0 mg/kg). In a within-subjects, counterbalanced design, mice received either vehicle + L-DOPA or M_5_ PAM VU0238429 + L-DOPA. Beginning 20 min after drug administration, animals were video recorded in a 3D Neurobehavioral Chamber (3D NC, see supplement) for one min every 20 min for 140 min. Videos were scored by a trained rater who was blind to condition using the validated rodent abnormal involuntary movements scale (AIMs) of L-DOPA-induced dyskinesia (LID). Data were analyzed for each time point using Wilcoxon Signed Ranked Test for each time point and for the overall sum of Axial, limb and orolingual (ALO) behavior. A) Shows the timeline of our experiments following initial motor testing on M_5_ modulators. B) Shows the median axial, limb, and orolingual (A.L.O.) sums for each time point for both vehicle + L-DOPA 1.5 mg/kg and M_5_ PAM, VU0238429 in black. The inset graph shows the overall sum of A.L.O. behaviors during the 140 min of ratings. AIMs data were analyzed by non-parametric Kruskal Wallis ANOVA with Dunn’s post-hoc tests employed as appropriate. Overall, there were no significant differences in overall A.L.O. behaviors, nor were there differences at any specific time point. C) A.L.O. AIMs data are shown for every 20 min for 140 min for Vehicle + L-DOPA 3.0 and M5 PAM + L-DOPA 3.0. The inset graph represents the overall sum of A.L.O. behaviors. Overall, there were no significant differences in overall A.L.O. sums nor were their differences in timecourse of A.L.O. behavior. D) Shows A.L.O. AIMs on vehicle + L-DOPA 6.0 or M5 PAM + L-DOPA 6.0. Again, mirroring other doses, there were no significant differences in overall A.L.O. AIMs scores nor in timecourse of A.L.O. behavior. E) We compared A.L.O. AIMs scores for peak plasma concentrations of M5 PAM alone or L-DOPA alone (1.5, 3.0, 6.0 mg/kg) using a Friedman ANOVA with test. Overall, we found that PAM did not produce robust dyskinesia, in contrast to L-DOPA alone. Data were analyzed using Friedman ANOVA with Dunn post-hoc tests as appropriate, * p < 0.05, *** p < 0.001

## Discussion

Because of the efficacy of non-selective anti-muscarinic compounds at reducing parkinsonian motor deficits, there has been a unique focus on understanding how blocking the signaling of acetylcholine signaling through specific muscarinic acetylcholine receptors(10, 14, 15, 17, 20, 29, 43). This work has indicated that M_1_ and especially M_4_ have a unique role in regulating the basal ganglia, and these receptors underlie the efficacy of non-selective anti-muscarinic therapeutics(13, 15, 16, 20, 29, 44). However, this may have hidden a role for the M_5_ muscarinic acetylcholine receptor in PD therapeutics. Elucidating the role of M_5_ muscarinic acetylcholine receptors has been challenging given the difficulty of creating a truly selective M_5_ receptor targeting compounds. However, discovery of the M_5_ PAM VU0238429 and M5 NAM ML375 which both have *in vivo* selectivity for M_5_ over the other muscarinic acetylcholine receptors have allowed for careful dissection of the role of M_5_ both in normal physiology and disease states (31, 32). Our current study, investigated the therapeutic potential of M_5_ receptor modulation in the unilateral 6-OHDA lesioned model of parkinsonian-like motor deficits and dyskinesia, and provides the first evidence that M_5_ is a viable target to provide anti-parkinsonian relief. M_5_ receptors have the potential for very targeted therapeutic benefit given that they are thought to be located exclusively on cell bodies of dopaminergic neurons in the substantia nigra pars compacta and on nigrostriatal DA terminals (12, 21, 22, 24). We found that M_5_ modulation could improve multiple aspects of motor performance in the 6-OHDA hemi-parkinsonian mouse. Specifically, we found that spatial aspects of gait, bradykinesia, and forepaw asymmetry were alleviated by M_5_ PAM administration alone. Furthermore, when combined with L-DOPA, M_5_ compounds do not negatively affect the motor benefit of L-DOPA, and suggest that L-DOPA and M_5_ may act through discrete pathways to exert anti-parkinsonian efficacy. Finally, we found that M_5_ receptor modulation does not potentiate L-DOPA-induced dyskinesia and does not independently cause dyskinesia. Taken together, these findings suggest that M_5_ PAM can be used to improve Parkinsonian gait deficits and bradykinesia without dyskinesia liability. This represents a highly novel new therapeutic pathway that may allow for anti-parkinsonian efficacy without adverse dyskinetic side effects. Previous work has shown that M_5_ NAMs lessen behaviors associated with drug abuse and craving(26–28, 45). Together with our data, this suggests potent modulation of DA neurons by M_5_ compounds, and may suggest a novel mechanism of action to block dopamine neuron firing or firing patterns in the treatment of dopamine associated diseases rather than just altering extracellular dopamine release, or its action on receptors. This may allow for modulation of DA signaling without using DA receptor targeting compounds which have broad effects on memory, cognition, motor behaviors, and impulse control disorders(46, 47), representing a novel class of anti-parkinsonian based therapeutics that may avert some adverse effects associated with L-DOPA or other dopaminergic medications.

Our data also highlights a novel use of a powerful, automated tool to phenotype motor movement in movement disorders models, the Erasmus ladder. This automated phenotyping tool captures multiple measures of motor performance, coordination, gait, and even motor learning. 6-OHDA lesioning impaired multiple measures of motor performance and coordination in line with what is expected from parkinsonian models. This tool allows for the automated, unbiased measurement of motor performance which could enhance rigor and reproducibility across studies given that many other motor behavioral tests are rated by observers and open to potential bias.

This study adds to a growing body of literature which explores selective modulation of muscarinic acetylcholine receptors as a novel therapeutic strategy to ameliorate Parkinson’s disease motor symptoms. Further studies will be required to understand how M_5_ modulation alters DA neuron firing and release across multiple models of PD to mechanistically understand how M_5_ exerts its observed efficacy here. One novel possibility is that M_5_ expression patterns are altered in response to loss of dopamine neurons, and that other neurons now express M_5_ other than midbrain dopaminergic neurons. Our data could be explained by new neuron types being modulated by M_5_ in addition to altering remaining DA neurons in this lesion model.

## Conclusions

M_5_ potentiation has the potential to remove parkinsonian motor deficits without a severe dyskinesia liability, and our data potentially point to efficacy in aspects of gait which are not well treated by current standard of care. Overall, our data highlight a completely novel therapeutic pathway to improve parkinsonian motor deficits without dyskinesia development or risk, and could represent a new therapeutic pathway to meet multiple unmet clinical needs in the management of PD.

## Author Contributions

**NEC:** Conceptualization, Project administration, Data curation, Formal Analysis, Visualization, Writing original draft, writing revision and editing

**DH:** Data curation, Writing revision and editing, approving final draft

**MK:** Data curation, writing revision and editing, approving final draft

**SG:** Data curation, writing revision and editing, approving final draft

**TE:** Data curation, editing revision, approving final draft

**MSM:** Funding acquisition, Project administration, Conceptualization, Formal Analysis, Writing revision and editing

**Data availability statement:** Data included in this manuscript will be available from the corresponding author by reasonable request.

## Financial Disclosure/Conflict of Interest

The Authors have no competing interests to disclose

## Funding

This work was supported by a target advancement award from the Michael J. Fox Foundation, MJFF-021787

## Notes

### Competing Interest Statement

The authors have declared no competing interest.

